# Loss of finger control complexity and intrusion of flexor biases are dissociable in finger individuation impairment after stroke

**DOI:** 10.1101/2023.08.29.555444

**Authors:** Jing Xu, Timothy Ma, Sapna Kumar, Kevin Olds, Jeremy Brown, Jacob Carducci, Alex Forrence, John Krakauer

## Abstract

The ability to control each finger independently is an essential component of human hand dexterity. A common observation of hand function impairment after stroke is the loss of this finger individuation ability, often referred to as enslavement, i.e., the unwanted coactivation of non-intended fingers in individuated finger movements. In the previous literature, this impairment has been attributed to several factors, such as the loss of corticospinal drive, an intrusion of flexor synergy due to upregulations of the subcortical pathways, and/or biomechanical constraints. These factors may or may not be mutually exclusive and are often difficult to tease apart. It has also been suggested, based on a prevailing impression, that the intrusion of flexor synergy appears to be an exaggerated pattern of the involuntary coactivations of task-irrelevant fingers seen in a healthy hand, often referred to as a flexor bias. Most previous studies, however, were based on assessments of enslavement in a single dimension (i.e., finger flexion/extension) that coincide with the flexor bias, making it difficult to tease apart the other aforementioned factors. Here, we set out to closely examine the nature of individuated finger control and finger coactivation patterns in all dimensions. Using a novel measurement device and a 3D finger-individuation paradigm, we aim to tease apart the contributions of lower biomechanical, subcortical constraints, and top-down cortical control to these patterns in both healthy and stroke hands. For the first time, we assessed all five fingers’ full capacity for individuation. Our results show that these patterns in the healthy and paretic hands present distinctly different shapes and magnitudes that are not influenced by biomechanical constraints. Those in the healthy hand presented larger angular distances that were dependent on top-down task goals, whereas those in the paretic hand presented larger Euclidean distances that arise from two dissociable factors: a loss of complexity in finger control and the dominance of an intrusion of flexor bias. These results suggest that finger individuation impairment after stroke is due to two dissociable factors: the loss of finger control complexity present in the healthy hand reflecting a top-down neural control strategy and an intrusion of flexor bias likely due to an upregulation of subcortical pathways. Our device and paradigm are demonstrated to be a promising tool to assess all aspects of the dexterous capacity of the hand.

## Introduction

Finger individuation, the ability to move our fingers independently, is of crucial importance to human hand dexterity. Loss of finger individuation after stroke is one of the hallmarks of upper extremity hemiparesis (Lang and Schieber 2003; Xu et al. 2017). A typical observation is that when a stroke survivor attempts to move one finger, undesirable coactivations from other fingers interfere with the intended finger action, a phenomenon referred to as finger enslavement. The nature of this impairment, however, remains to be elucidated. Some possible contributors have been suggested in the literature, such as the loss of complexity and flexibility, i.e., the large repertoire, in finger control due to damage of corticospinal pathways (Lang and Schieber 2004a; Xu et al. 2017), an intrusion of flexor synergy (Ellis, Schut, and Dewald 2017) due to upregulation of subcortical pathways, and possible unchecked biomechanical constraints after loss of neural control.

While these factors may or may not share the same underlying mechanisms, it is difficult to tease apart their roles in finger individuation, partly due to limitations in measurement techniques to accurately assess the full capacity of finger control. Previous work has largely focused on finger flexion, mainly involving one- or two-dimension finger flexion-extension at the metacarpophalangeal (MCP) joints. For example, assessed by a keyboard-like device to examine finger flexion, enslaving patterns in the paretic hand in our earlier study appeared to present an overall scaling up of magnitude but a similar shape compared to the non-paretic hand (Xu et al. 2017; Ejaz et al. 2018). Likewise, finger individuation studies using an instrumented glove, the Cyberglove (Virtual Technologies, Palo Alto, CA), predominately examined flexion/extension movements, because measures in the ab/adduction directions using the Cyberglove are not reliable (Lang and Schieber 2004b). The coactivation patterns observed in these tasks largely coincide with a natural flexor inclination, as the human hand is evolved to possess strong prehension abilities (Napier 1956), leading to an impression that enslavement in stroke patients is an exaggerated version of the prehension bias seen in the healthy hand (Z. M. Li, Latash, and Zatsiorsky 1998; S. Li et al. 2003; Abolins and Latash 2021). However, this impression may be largely influenced by the limited capacity to capture the full range of finger control.

A few studies that did test other movement directions in stroke-affected hands suggest differential impairment of finger independence in flexion, extension, and ad/abduction (Lang and Schieber 2003; 2004a). Moreover, studies that assessed naturalistic movements of the hand in healthy people have also shown that finger coactivation patterns can vary greatly depending on the tasks and fingers involved (Ingram et al. 2008; Lang and Schieber 2004b; Todorov and Ghahramani 2004; Yan et al. 2020), and that training finger individuation in the flexion direction does not transfer to the extension direction (Kamara et al. 2023). Our recent finger flexion study (Xu et al. 2017) suggests that finger individuation and strength follow different recovery processes and that they may be supported by different neural pathways, with individuation mainly supported by the corticospinal tract (CST) and strength largely driven by the reticulospinal tract (RST). Thus, there seems to be a logical gap when assuming that finger enslaving patterns due to the loss of CST after stroke would present a similar shape as coactivation patterns in a healthy system.

These findings raise the question as to what extent the enslaving patterns after stroke resemble the coactivation patterns observed in the healthy hand and, subsequently, whether they share the same biomechanical and neural sources. Do they differ only in *magnitude* or also in *shape* when it comes to movements in directions outside the prehension synergy? Which ones among the three factors: complexity, intrusion of flexor bias, and unchecked biomechanical constraint, dominate the finger individuation impairment after stroke?

Here we set out to examine the full extent of finger individuation impairment after stroke and tease apart its possible underlying origins. We hypothesize that the three factors can be dissociable in finger individuation impairment, and that they are combined to contribute to the enslaving patterns in the paretic hand. We further hypothesize that the paretic enslaving patterns are neither an exaggerated pattern of the involuntary activity in task-irrelevant fingers, nor the prehension bias in the healthy hand. These coactivation patterns in the healthy hand, sometimes undesired but often out of convenience of striking a balance between precision and efficiency, may be qualitatively and mechanistically different from those observed in patients with diminished motor repertoire. To avoid conceptual confusion, we hereupon use the term *coactivation* to refer to the patterns observed in a healthy hand and *enslavement* to those in a paretic hand.

To accurately quantify finger activities in three dimensions (3D), we engineered a highly sensitive and ergonomic measurement device that allows simultaneous recording of small isometric forces at all five fingertips in 3D, the Hand Articulation Neuro-training Device (Carducci et al. 2022) (HAND, JHU reference #C14603, Fig. 1A-B). We then designed a 3D finger-individuation task (Fig. 2) to obtain a more complete picture of individuation ability and finger coactivation and enslavement patterns in healthy and paretic hands, respectively.

**Figure 1.**
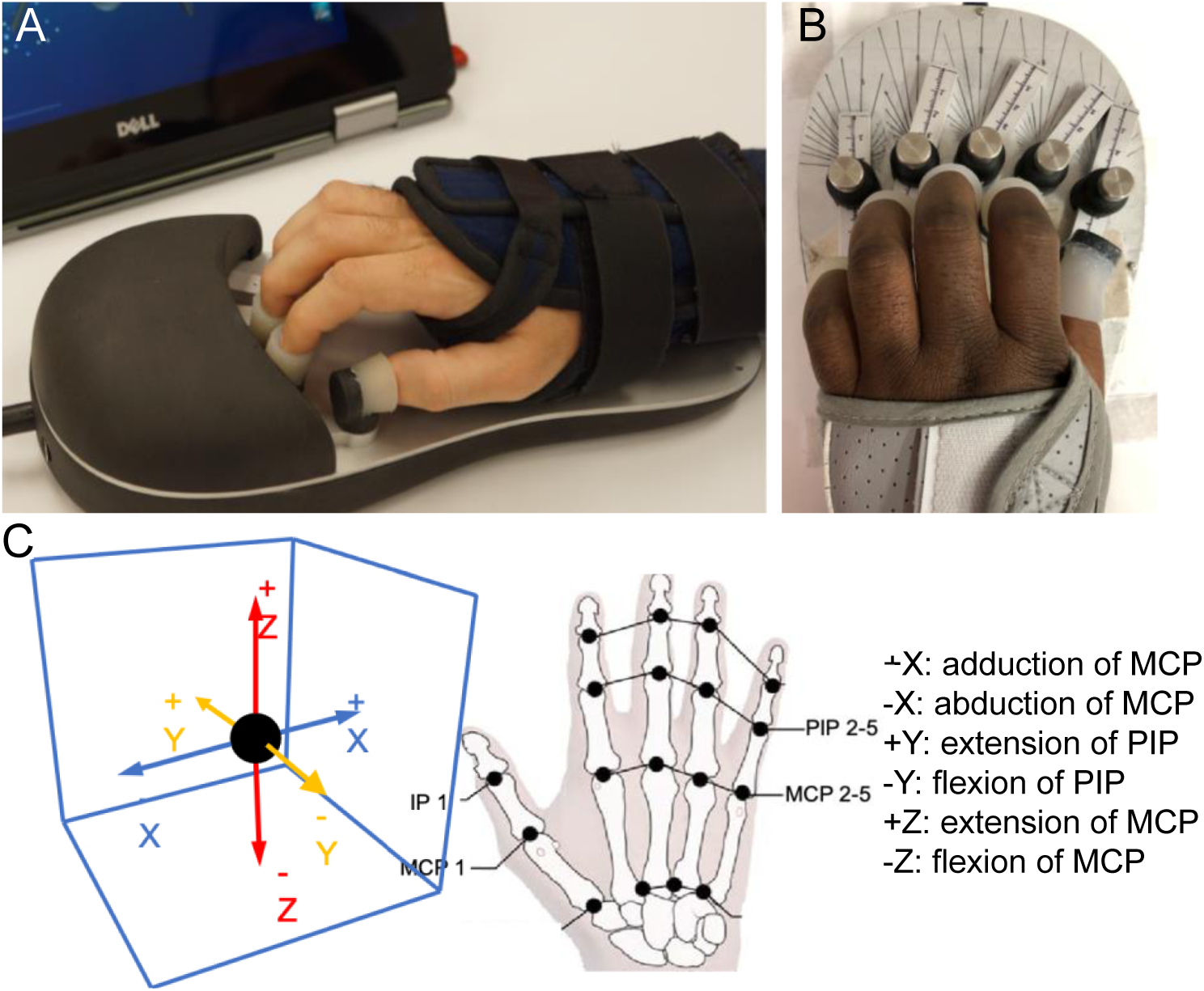
Hand Articulating Neuro-Training Device (HAND). **A-B**. Participants’ hand fitting in the device, posture recorded with mounting distance and angle. **C**. Illustration of finger joint motion to Cartesian coordinates in the virtual 3D space.

**Figure 2.**
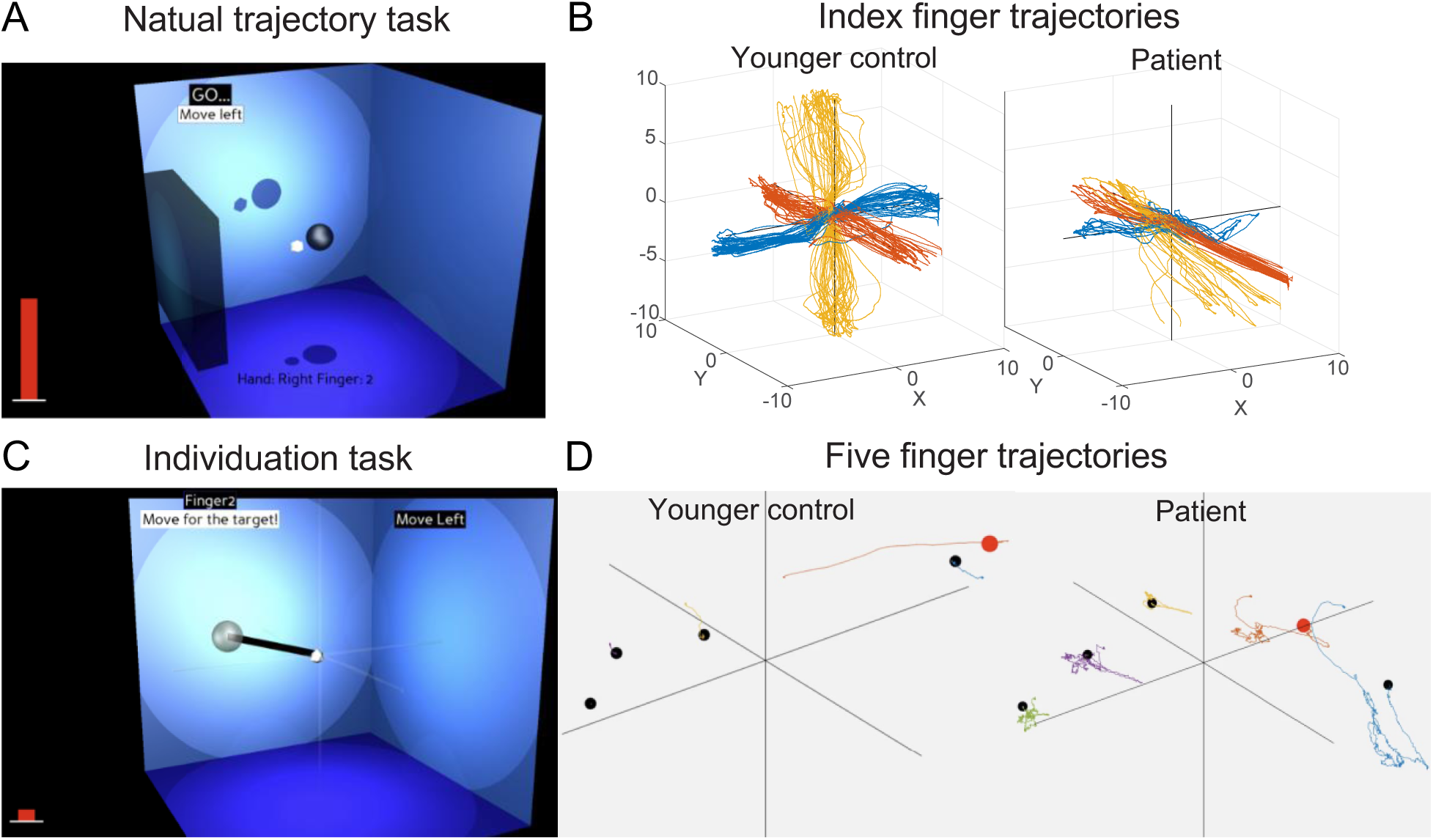
The 3D finger individuation paradigm. **A-B**. Natural trajectory task. **A.** Screenshot of the task for one finger: participant control a white dot by exerting isometric forces towards one of the 6 directions in the 3D virtual space and move the dot between the home position (gray sphere) and a virtual wall. **B.** Example force traces recorded from index finger during natural trajectory task in a healthy participant and a stroke patient’s paretic hand; **C-D**. Finger individuation task. **C.** Screenshot of the task for one finger: participant control a white dot in the virtual space and try to hit a target by following the specified path (thick black line) estimated from that finger’s natural trajectory (shown in B) while attempting to minimize forces from uninstructed fingers (red bar). D. Example force traces recorded from all five fingers in a healthy participant’s left hand and a stroke survivor’s paretic hand during the individuation task when the participants attempted to move their left index finger towards a target (red dot).

Due to its high-dimensional nature, hand behavior kinematics/kinetics analysis in 3D is challenging. Much of the prior work has adopted various dimensionality reduction approaches. To measure finger individuation, for example, an Individuation Index was derived from task-irrelevant finger activation levels as a function of activity in the instructed finger (e.g. Häger-Ross and Schieber, 2000; Xu et al., 2017). Another approach is variants of matrix factorization methods, such as principal component analysis (PCA) (e.g., (Santello, Flanders, and Soechting 1998)), nonnegative matrix factorization (NMF) (e.g., (Lee and Seung 1999)), or independent component analysis (ICA) (e.g., (Tresch, Cheung, and d’Avella 2006)). While these approaches have demonstrated considerable explanatory power, they involve substantial dimensional reduction, and thus lose the rich information of finger-specific coactivation patterns, especially the movement direction of each enslaved finger. Moreover, in synergy analyses, the modules are often difficult to interpret and vary greatly from task to task (Todorov and Ghahramani 2004). To analyze the 3D finger trajectories recorded from all five fingertips without losing the fine-grained details of finger- and direction-specific coactivation, we derived three measures (see details in Methods): 1) a 3D *Enslavement* metric as ratios of uninstructed/instructed finger forces, 2) a *Bias* metric as ratios of positive/negative direction forces for each active finger that captures each instructed finger’s tendency towards its preferred direction in voluntary movement, and 3) hand resting posture, the Mount Angle and Mount Distance where a hand was fit in the device that produced minimum force. We then used the Representational Similarity Analysis (RSA) approach (Kriegeskorte 2011; Kriegeskorte and Kievit 2013) and linear-mixed effect (LME) models to assess both magnitude and shape differences in these patterns, aiming to tease apart the three factors, complexity, flexor intrusion, and biomechanical constraints on coactivation/enslaving patterns. In RSA analysis, angular (cosine) distance is sensitive to the *directional* differences between patterns, whereas Euclidean distances are sensitive to the *length* difference between two force vectors but insensitive to direction differences (Walther et al. 2016). A larger difference in *direction* indicates qualitatively different *shapes* of coactivation patterns, whereas a larger difference in *length* indicates quantitatively different *magnitudes* of coactivation patterns. We thus assess the complexity of finger control by Angular Distance of finger coactivation/enslavement patterns, a gauge of difference in geometric *shapes*, when instructed fingers were engaged in different tasks; we use Euclidean Distance to assess the *magnitude* of the pattern differences across tasks. Flexor intrusion was assessed by Bias differences between the paretic and non-paretic hands in the flexion-extension dimensions, whereas biomechanical constraints were assessed by resting hand postures.

We operationally define the *top-down* factors in our task as *task goals*, specifically, instructed finger and target direction. If the finger coactivation patterns are driven by top-down factors, different instructed fingers and target directions will lead to different geometric *shapes* of the patterns, i.e., uninstructed finger patterns will follow the instructed finger depending on which finger is the primary mover and which direction it intends to move in. In contrast, if the patterns present similar shapes regardless of task goals, top-down cortical influence would be weak, implying a greater influence by lower-level biomechanical and/or subcortical neural constraints. To further tease apart the biomechanical vs. subcortical constraints that contribute to the enslavement pattern in the paretic hand, we used the resting hand postures and the healthy bias pattern, a flexor bias presumably dominated by biomechanical constraints, to predict the paretic bias and enslavement patterns. We then examined the extent to which flexor intrusion can account for the shape and magnitude of finger coactivation/enslavement patterns. We predicted that 1) compared to the healthy hand, the enslavement patterns in the paretic hand would present reduced complexity in geometric shapes, thus smaller angular distances across different task goals, but an increased magnitude, i.e., Euclidean distances; 2) Impairment of individuation and the increased magnitude of the enslaving patterns in the paretic hand would be dominated by the intrusion of flexor bias; 3) Intrusion of flexor bias, however, would not be predictive to the loss of complexity, i.e., angular distances in the paretic hand; Lastly, 4) none of the patterns in the healthy or paretic hands could be accounted for by biomechanical factors.

## Results

We tested 13 chronic stroke patients and 29 healthy participants’ finger individuation ability in 3D, while recording isometric fingertip forces from all five fingers simultaneously. Patient characteristics and clinical assessment results are reported in Table 1.

**Table 1.**
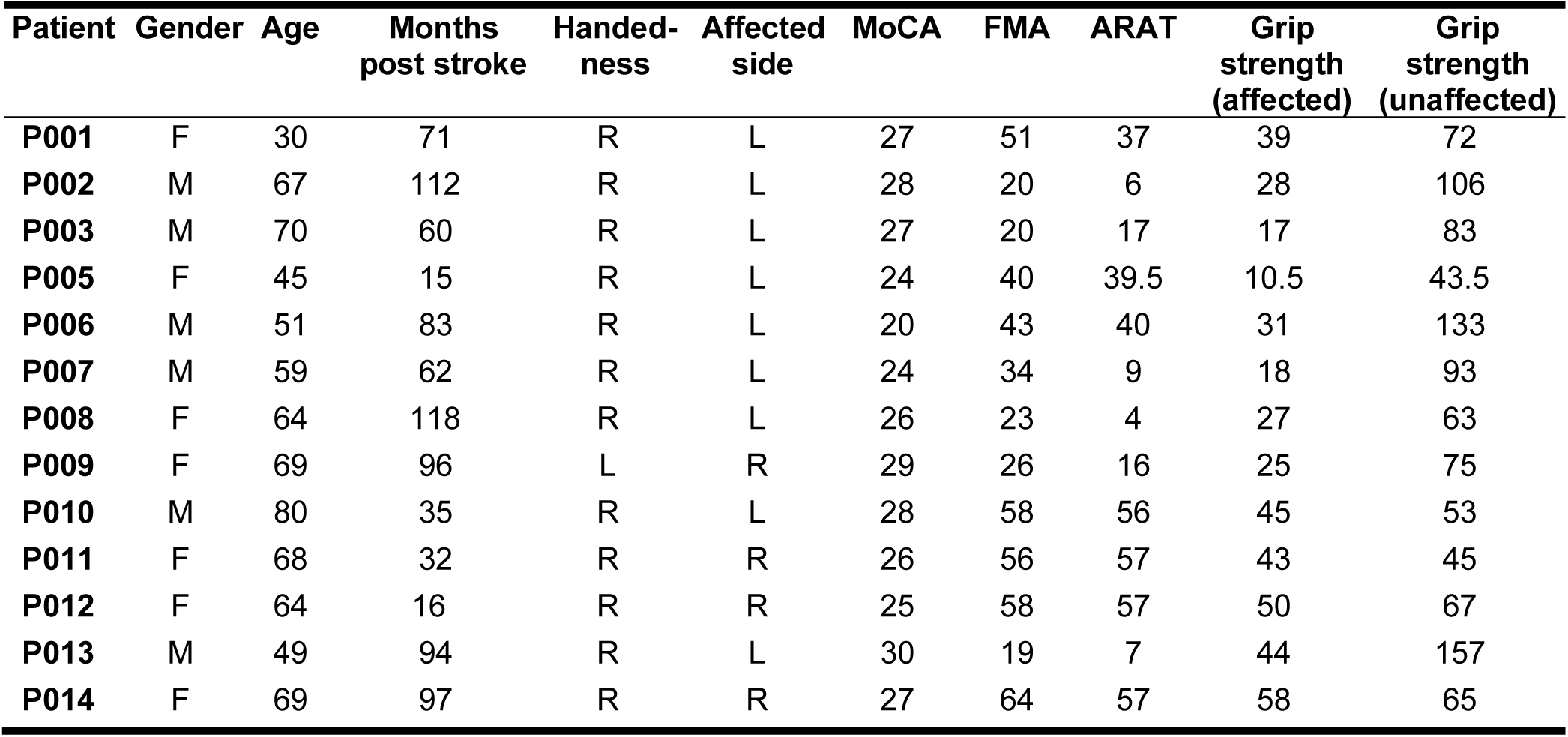
Patient characteristics. Data indicate patients’ gender (M, male; F, female), age (years), time since stroke (in months), handedness (L, left; R, right), affected side (L, left; R, right), Montreal Cognitive Assessment (MoCA, maximum 30), Fugl-Meyer Assessment for upper extremity (FMA, maximum 66), and Action Research Arm Test (ARAT), and grip strength (in pounds) if applicable.

High-resolution isometric forces were recorded from all five fingertips in 3D while a participant was engaged in a task of isolating one finger joint to move a dot reflecting their fingertip force projected on the computer monitor in a virtual 3D task space. Three major outcome measures, Individuation Index, Bias, and Enslavement, showed high reliability using the split-half reliability measure (see Materials and Methods): Individuation Index reliabilities were 0.98, 0.97, and 0.96 for healthy, non-paretic, and paretic hands, respectively; Bias reliabilities were 0.89, 0.85, and 0.90, for healthy, non-paretic, and paretic hands, respectively; Enslavement reliabilities were 0.85, 0.87, 0.82 for healthy, non-paretic, and paretic hands, respectively. We first report 3D Individuation Indices for each finger in both Cartesian and joint spaces. We then present results from three tests for our main hypotheses in the following sections. Lastly, we report results from PCA analysis where we compared the number of PCs required to account for variances in the healthy, non-paretic, and paretic hands’ 3D isometric forces in the individuation task.

### Finger individuation in paretic hand was differentially impaired across fingers and joints

To capture participants’ finger individuation abilities, we derived an Individuation Index, quantified as the slope of the linear function of the mean activation from uninstructed fingers against forces produced in the instructed finger (Xu et al. 2017) (Fig. 3A). In the current analysis, Individuation Index is calculated for six target directions in the 3D Cartesian space: +/- X, +/- Y, +/- Z (Fig. 1C), instead of a single direction (finger flexion). To gain insight into finger individuation abilities at the anatomical level and to connect our results to clinical knowledge, we mapped the six directions in the Cartesian space to the joint space (Fig. 1C).

**Figure 3.**
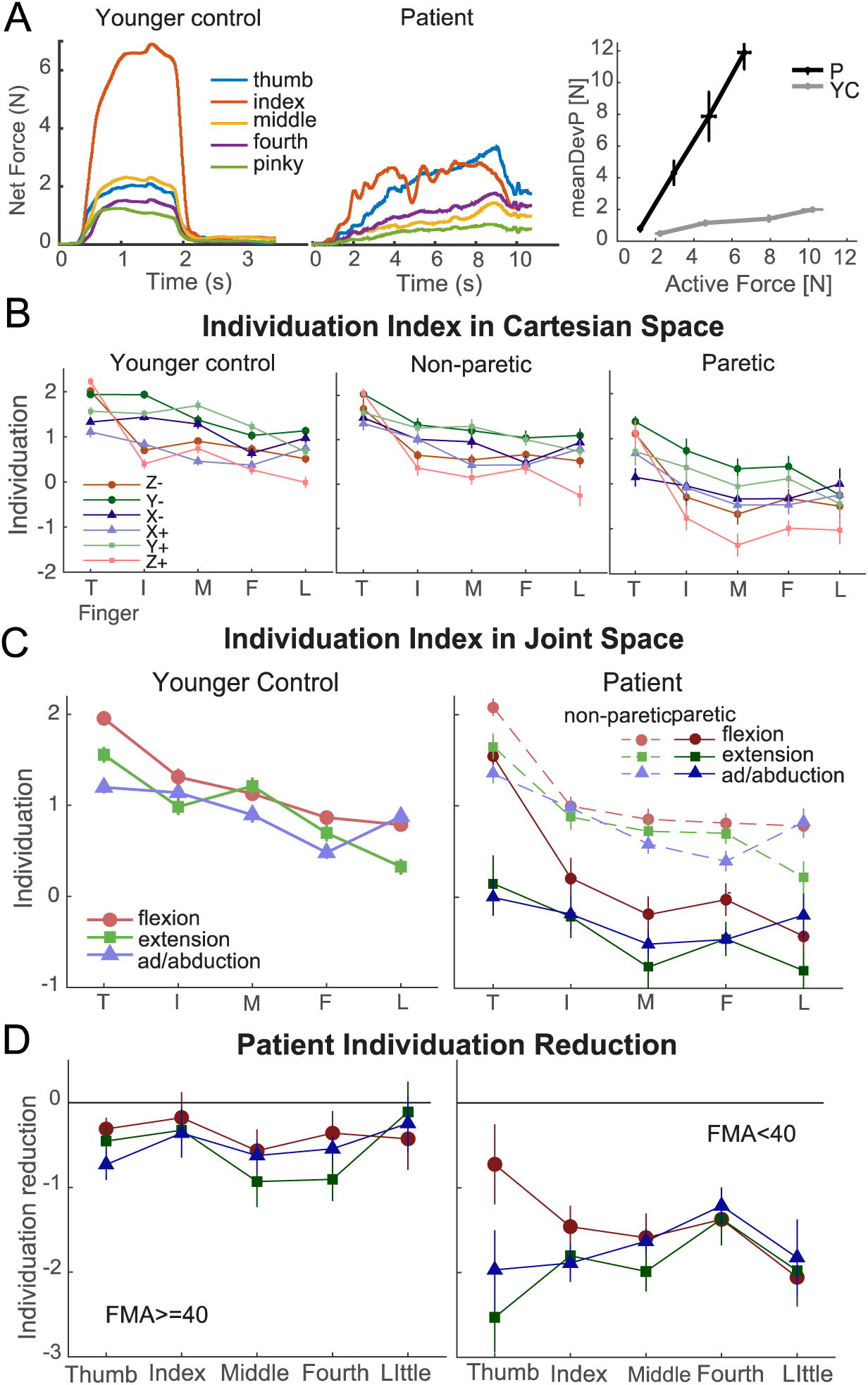
Individuation Index derivation and results. **A**. Illustration of derivation of Individuation Index: overall net force trajectories (first two panels) and the function of mean deviation net force from the uninstructed fingers as a function of the force in the instructed finger towards the instructed direction (3^rd^ panel). Individuation index is -log(slope) of the regression line of the function; **B**. Individuation Indices of healthy and stroke patients in cartesian spaces; **C**. Individuation Indices summarized in joint space for healthy controls and patients; **D.** Reduction of Individuation Indices (paretic subtracted from non-paretic) for patients with mild (FMA>=40) and severe (FMA<40) impairment.

Consistent with previous findings (Häger-Ross and Schieber 2000), our results showed that in younger healthy participants, there was a significant difference in Individuation Index across different fingers (likelihood ratio test against the null model for Instructed Finger effect *χ*^2^ = 67.77, *p* < 2.2e-16) with the thumb having the highest level of individuation (pair-wise comparison between thumb and other fingers: t(28) > 4.27, *p* < 0.0005) and the fourth- and the little-finger Individuation Indexes the lowest among all fingers, with no statistical difference between the two (*t*(28) = 0.09, *p* = 0.93) (Fig. 3B). Interestingly, finger flexion appeared to be the easiest direction to individuate across all fingers (pair-wise comparison between flexion and extension/adduction/adduction: *t*(28) > 8.38, *p* < 4.82e-7).

In stroke patients, the non-paretic hands presented a very similar trend across fingers/directions as younger healthy participants (Group effect not significant: *χ*^2^ = 0.10, *p* = 0.76) (Fig. 3B-C). In contrast, the paretic hand showed a clear reduction of individuation ability across all fingers and all directions compared to the non-paretic hand (Group effect *χ*^2^ = 70.10, *p* < 2.2e-16). Movement in the joint space (flexion/extension/abduction/adduction) had a significant interaction with the Instructed Finger (*χ*^2^ = 14.27, *p* = 0.00016), indicating that different movement directions across different fingers were impaired differentially. Interestingly, finger extension and abduction/adduction were more impaired than flexion (pair-wise comparison between flexion and extension: *t*(12) = 2.89, *p* = 0.014; flexion and abduction/adduction: *t*(12) = 2.83, *p* = 0.015), but impairment of extension and abduction/adduction were not significantly different (*t*(12) = −1.35, *p* = 0.20) (Fig. 3C). This finding is consistent with clinical knowledge and previous findings (Lang and Schieber 2004a).

To examine the impact of stroke severity on impairment of finger individuation, we further sub-divided patients by a standard clinical assessment, Fugl-Meyer Upper Extremity Assessment (FMA), to mild (FMA ≥ 40, N=7) vs. moderate-severe (FMA < 40, N=6) groups. As shown in Fig. 3D, the two groups of patients present very different patterns of individuation impairment: while the overall impairment in mild patients shows a similar level of reduction from non-paretic hand across all fingers except for the little finger, moderate-severe patients show a quite different level of impairment across fingers and movement directions: thumb flexion for these patients are the least impaired and the only movement direction close to those in the mild or healthy hand; thumb ad/abduction and little-finger flexion were the most impaired. For both groups of patients, the impairment of the thumb and little finger showed opposite patterns (Fig. 3C-D): the least impaired direction for the thumb was flexion, whereas that for the little finger was extension. These distinct patterns across impairment severity and fingers/target directions suggest that there may be different finger enslavement patterns underneath Individuation Indexes.

The Individuation Index, a gross summary across all finger activities, does not contain important information regarding the direction of coactivation in the uninstructed fingers, because the 3D forces in all uninstructed fingers were summed to a net force regardless of their directions in the 3D space. Therefore, while informative, the apparent DC-shift of Individuation Index, i.e., reduction of magnitude with a preserved overall shape, from the non-paretic to paretic hand at first glance (Fig. 3C) could be misleading, giving the impression that the enslaving pattern observed after stroke only differs in *magnitude* from the healthy pattern. To examine the *shape* of finger coactivation patterns in the healthy and stroke hand, we need to go beyond the Individuation Index. In the following sections, we report these shape differences using RSA and LME analyses and relate these analyses to a lower-level finger bias measure to examine three possible contributing factors to the impairment of finger individuation: loss of complexity, intrusion of flexor bias, and biomechanical constraints.

### Finger coactivation patterns in the paretic hand present reduced shape variance across task goals

We tested the hypothesis that the complexity, i.e., geometric shapes, of finger coactivation patterns with the uninstructed fingers across task goals will be reduced after stroke, whereas these patterns in the healthy and non-paretic hands would be mainly driven by top-down task goals, i.e., which finger to move (Instructed Finger) and where to move to (Target Direction), and would present more variable shapes across task goals. If task goals have a major influence on the activities observed in uninstructed fingers, these patterns will be different in *shape* depending on the primary finger and target direction, whereas if the coactivation patterns are mainly driven by low-level constraints, they may only differ in *magnitude* regardless of instructed finger and target direction.

To directly assess shape vs. magnitude differences across finger coactivation/enslaving patterns, we used Angular and Euclidean Distance measures between finger coactivation patterns across instructed fingers and target directions. While Angular Distance is more sensitive to the directional, hence shape, differences between finger coactivation patterns across different task goals, Euclidean Distance is more sensitive to the length, hence magnitude, differences. Representation distance matrices (RDMs) were constructed for each Instructed Finger and Target Direction for both distance measures by computing the pairwise distances between uninstructed finger coactivation patterns across two Target Directions for each Instructed Finger or between different Instructed Fingers for each Target Direction (Fig. 4A, D, see Materials and Methods). The mean distances from these RDMs were then compared across paretic and non-paretic hand using a pair-wise t-test.

**Figure 4.**
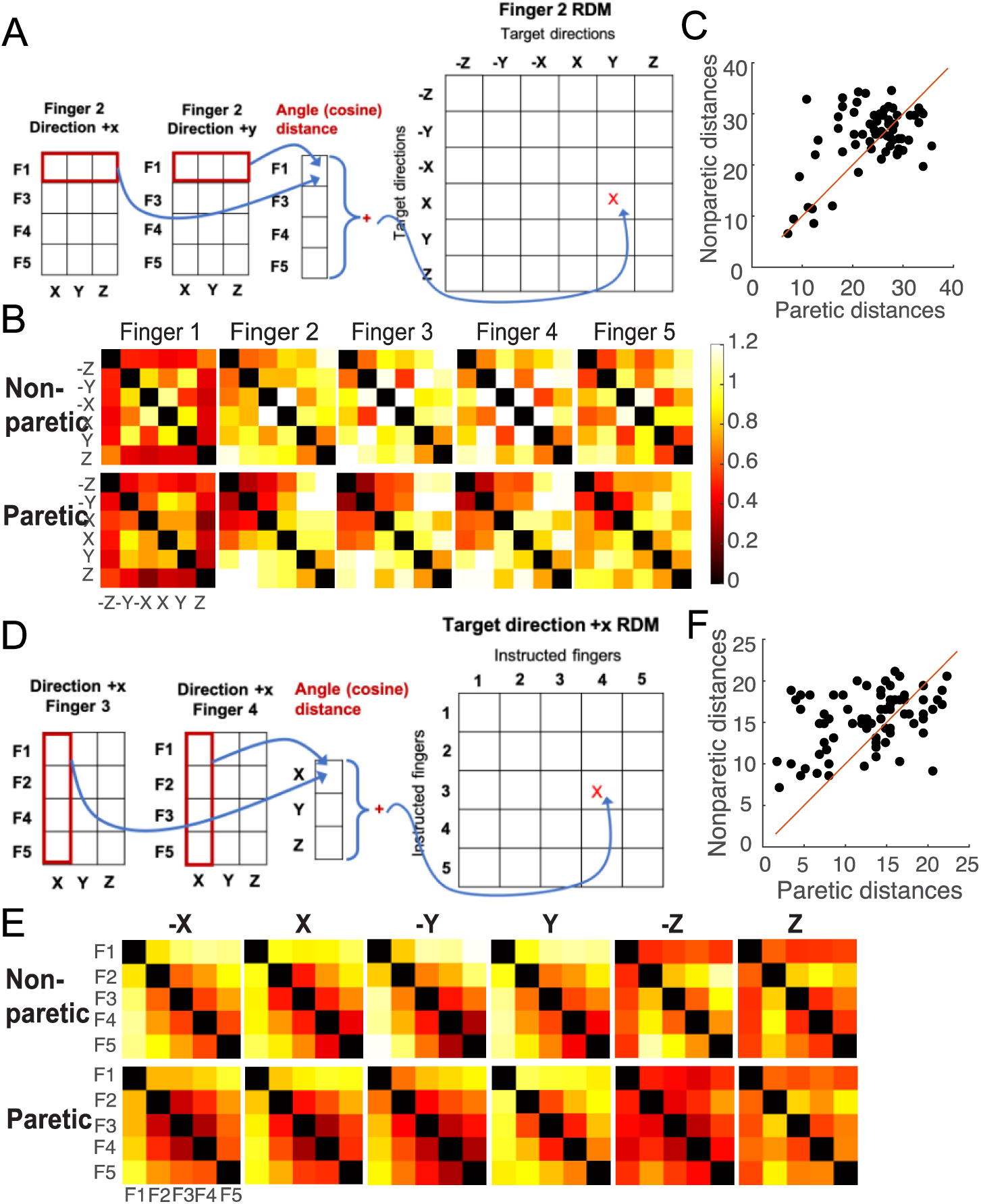
Pattern similarity analysis of finger coactivation/enslavement patterns using cosine distances. A larger distance indicates distinct shapes of finger coactivation patterns. **A.** An illustration of representation similarity matrix (RDM) computed across coactivation/enslavement patterns for one instructed finger (finger 2) exerting force in two different directions: +X vs. +Y. **B.** Mean RDMs for each instructed finger direction averaged across all non-paretic and paretic hands. **C.** Direct comparison of mean distance values for each participant at each instructed finger. **D.** An illustration of RDM computed across coactivation/enslavement patterns for two different fingers (finger 3 & 4) exerting force in the same target direction (+X). **E.** Mean RDMs for each target direction averaged across all non-paretic and paretic hands. **F.** Direct comparison of mean distance values for each participant at each instructed target direction.

The results support our hypothesis that the paretic hand would show reduced shape variance and, thus, complexity across different task goals. Visual inspection of all mean RDMs across all patients showed consistently higher Angular Distances in the non-paretic hand than in the paretic hand (Fig. 4B, E). Pair-wise t-tests across the paretic and non-paretic hands for comparisons across Instructed Fingers (*t*(64) = −1.92, *p* = 0.059) and Target Directions (*t*(77) = - 4.42, *p* = 3.24e-05) also confirmed this visual impression (Fig. 4C, F). There is a gradient of reduction of Angular Distances from mild to more severely impaired patients (Fig. 5) (Independent t-tests between mild and severe patients: by Instructed Finger *t*(11) = 4.39, *p* = 0.001; by Target Direction *t*(11) = 1.16, *p* = 0.27). Interestingly, pattern distances by Target Direction, i.e., between different fingers moving towards the same directions, were only significantly different between mild and severe patients for directions -X, -Y & -Z (*t*(11) = 3.20, *p* = 0.008), indicating a reduction of Angular Distances in more severe patients in the flexion movement directions.

**Figure 5.**
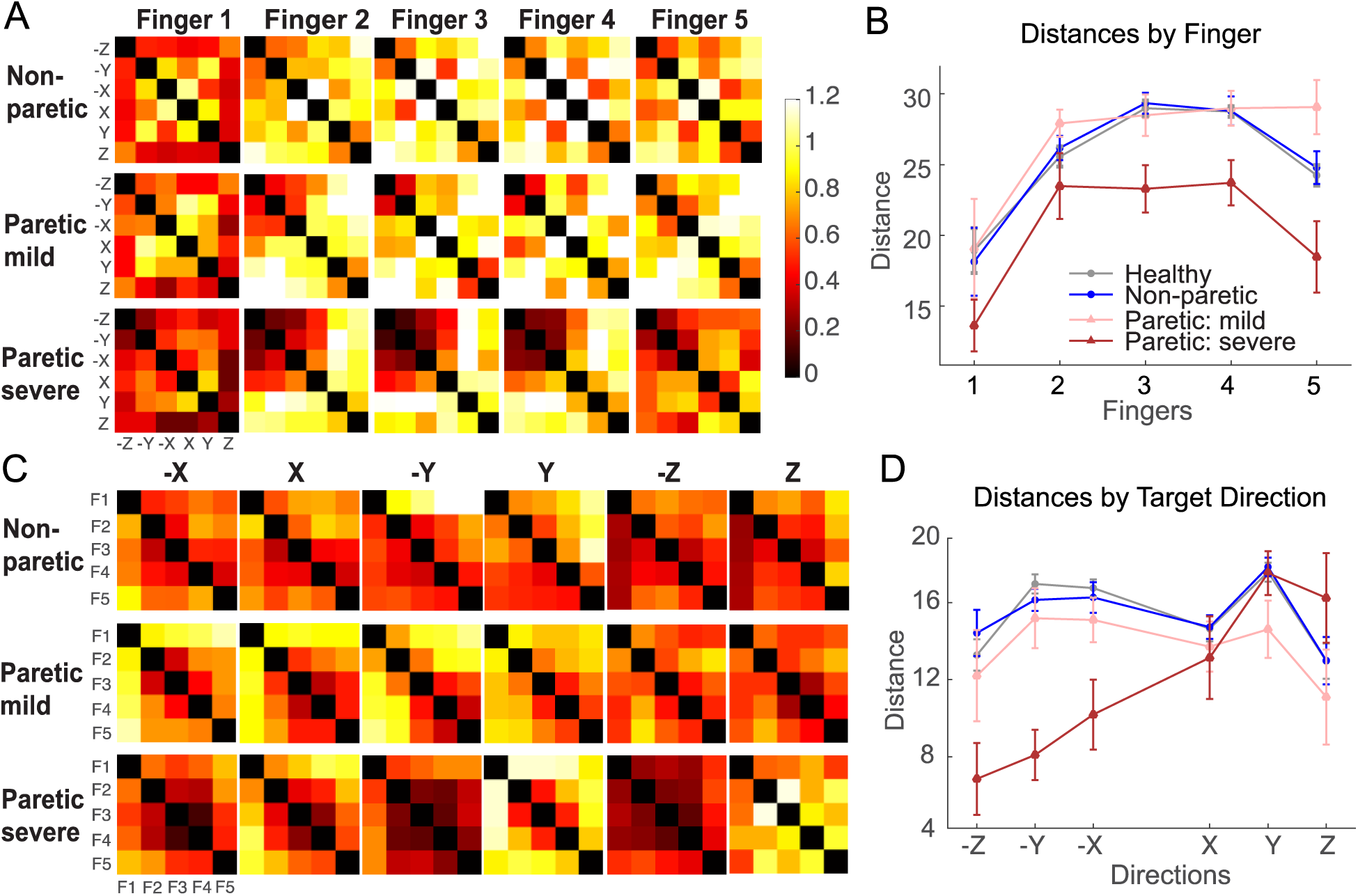
Pattern similarity analysis of the finger coactivation/enslavement patterns using Cosine distances with mildly (FMA>=40) and severely (FMA<40) impaired patients separated. **A.** Mean RDMs for each instructed finger direction averaged across all non-paretic and paretic hands. **B.** Sum of mean distance values across all participants for each instructed finger. **C.** Mean RDMs for each target direction averaged across all non-paretic and paretic hands. **D.** Sum of mean distance values across all participants for each instructed target direction.

Euclidean Distances showed an opposite pattern of results: significantly larger distances in the paretic hand compared to the non-paretic hand (By Instructed Finger *t*(64) = 8.75, *p* = 1.55e-12, by Target Direction *t*(77) = 7.78, *p* = 2.74e-11, Fig. 6). This finding indicates that the paretic hands showed enslavement patterns in similar shapes that differ only in magnitude across different task goals (i.e., Instructed Finger and Target Direction). Moreover, in contrast to Angular Distances, more severely impaired patients showed a stronger magnitude of Euclidean Distances across enslaving patterns for different task goals (Independent t-tests: Target Direction *t*(11) = −3.92, *p* = 0.002, Instructed Finger *t*(11) = −3.78, *p* = 0.003) (Fig. 7).

**Figure 6.**
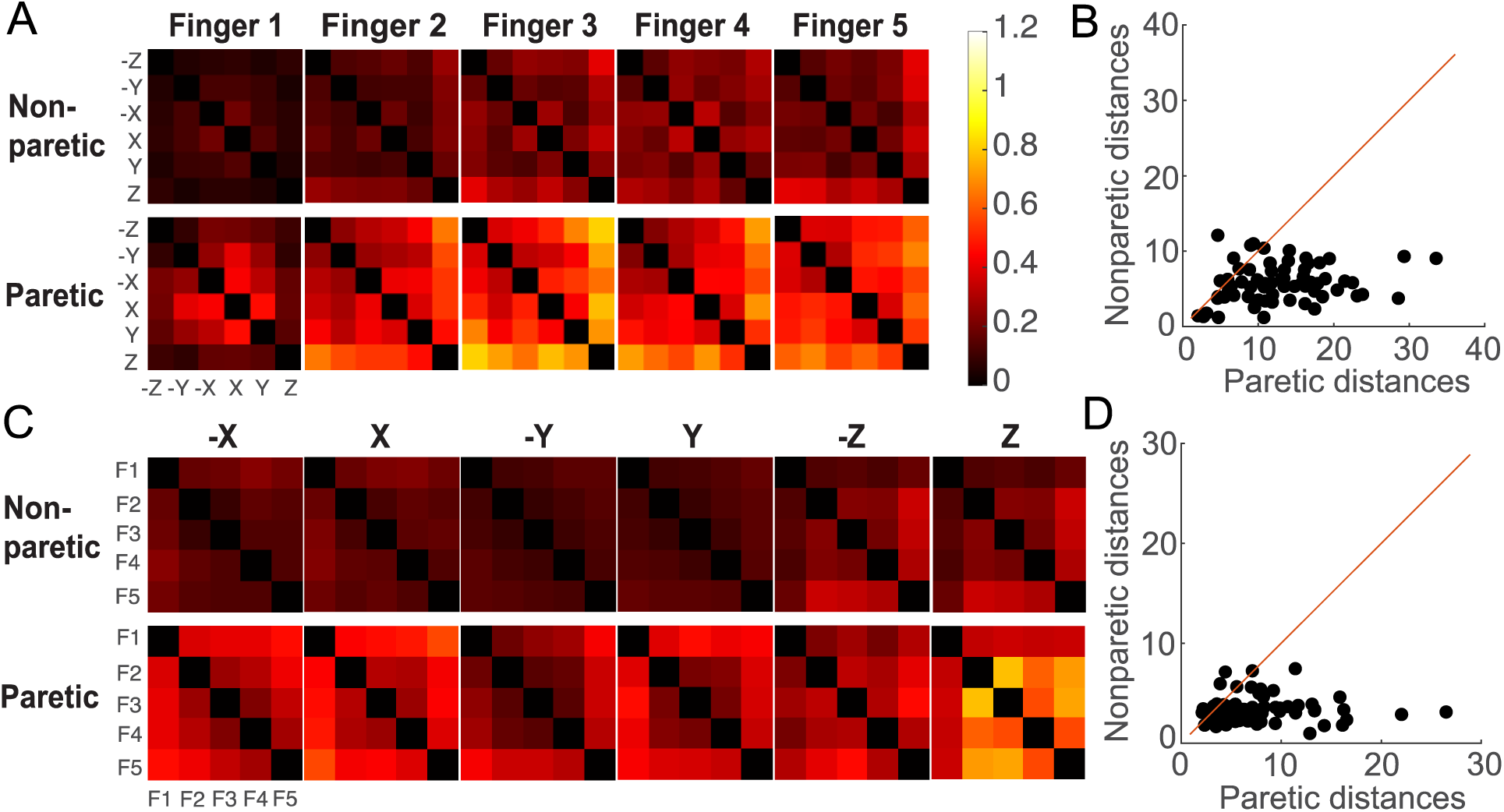
Pattern similarity analysis of the finger coactivation/enslavement patterns using Euclidean distances. **A**. Mean RDMs for each instructed finger direction averaged across all non-paretic and paretic hands. **B**. Direct comparison of mean distance values for each participant at each instructed finger. **C**. Mean RDMs for each target direction averaged across all non-paretic and paretic hands. **D**. Direct comparison of mean distance values for each participant at each instructed target direction.

**Figure 7.**
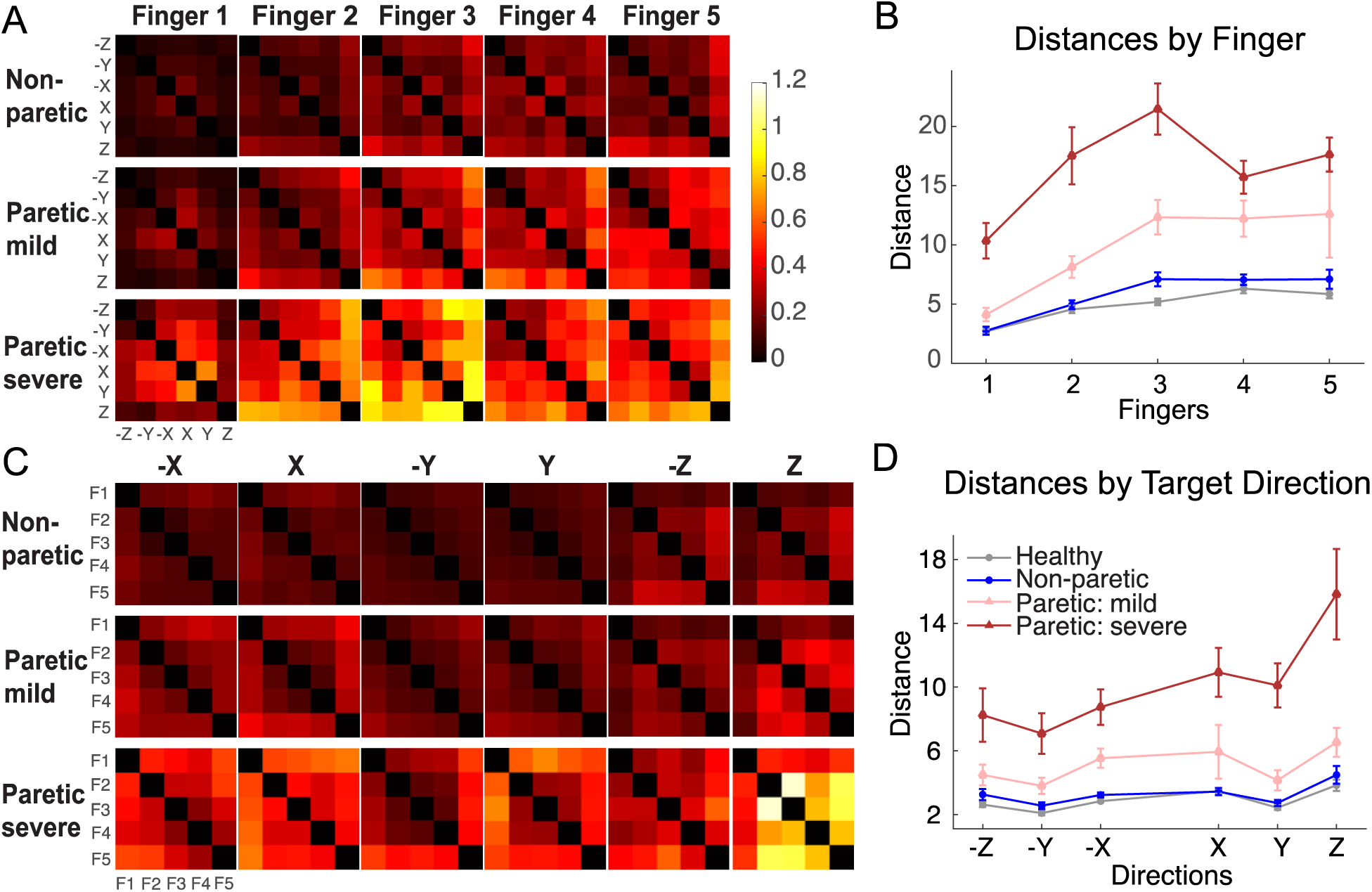
Pattern similarity analysis of finger coactivation/enslavement patterns using Euclidean distances, with mildly (FMA>=40) and severely (FMA<40) impaired patients separated. **A.** Mean RDMs for each instructed finger direction averaged across all non-paretic and paretic hands. **B.** Sum of mean distance values across all participants for each instructed finger. **C.** Mean RDMs for each target direction averaged across all non-paretic and paretic hands. **D.** Sum of mean distance values across all participants for each instructed target direction.

### Subcortical but not biomechanical constraints dominate enslavement patterns in the paretic hand

In this section, we explore the sources of the increased magnitude of enslaving patterns after stroke. Specifically, we tested the effect of top-down (task goals), low-level subcortical (biases), and biomechanical factors (hand resting postures) on finger coactivation/enslavement patterns and Individuation index in the paretic and non-paretic hands.

We used two measures to fully account for the low-level (biomechanical and/or subcortical) factors that may contribute to the finger coactivation/enslavement patterns (see details in Materials and Methods): 1) a *bias* measure, a log ratio of the active finger forces along two directions of each Cartesian axis, to quantify each finger’s natural tendency within its range of motion, which presumably captures the active/passive imbalance between agonist and antagonist muscles that can be driven by subcortical and biomechanical sources (Fig. 1C); and 2) the resting hand postures, measured by *Mount Angle* and *Mount Distance* when the hand is fit to the device with minimum force recorded (Fig. 1B), which are presumably dominated by passive biomechanical features.

Visual inspection of the overall patterns of Biases plotted in the 3D space suggests that the healthy, non-paretic, and paretic hands show very similar trends with larger values in the -X, -Y, and -Z directions, reflecting a flexor bias (Fig. 8A). The paretic hand Biases have larger amplitudes. LME model with Instructed Finger, Target Direction, Group (healthy, non-paretic), Mount Distance, and Mount Angle showed that Biases were not significantly different between the healthy and the non-paretic hands (Group effect *p* = 0.31). In contrast, biases in the paretic hand were significantly higher compared to those in the non-paretic hand (*p* = 4.0e-11). In the analysis thereinafter, *non-pathological hands* will be used to refer to the healthy and non-paretic hands combined in one group, and *non-paretic hands* will be used to refer to the unaffected side of the stroke participants. In all groups, a flexor bias is present, indicated by a higher magnitude in -Y and -Z directions (non-pathological hands -Y: *t*(44) = 5.19, *p* = 2.58e-6, -Z: *t*(44) = 9.64, *p* = 1.01e-12; paretic hands -Y: *t*(12) = 3.25, *p* = 0.004, -Z: *t*(12) = 4.12, *p* = 0.0007) (Fig. 8B).

**Figure 8.**
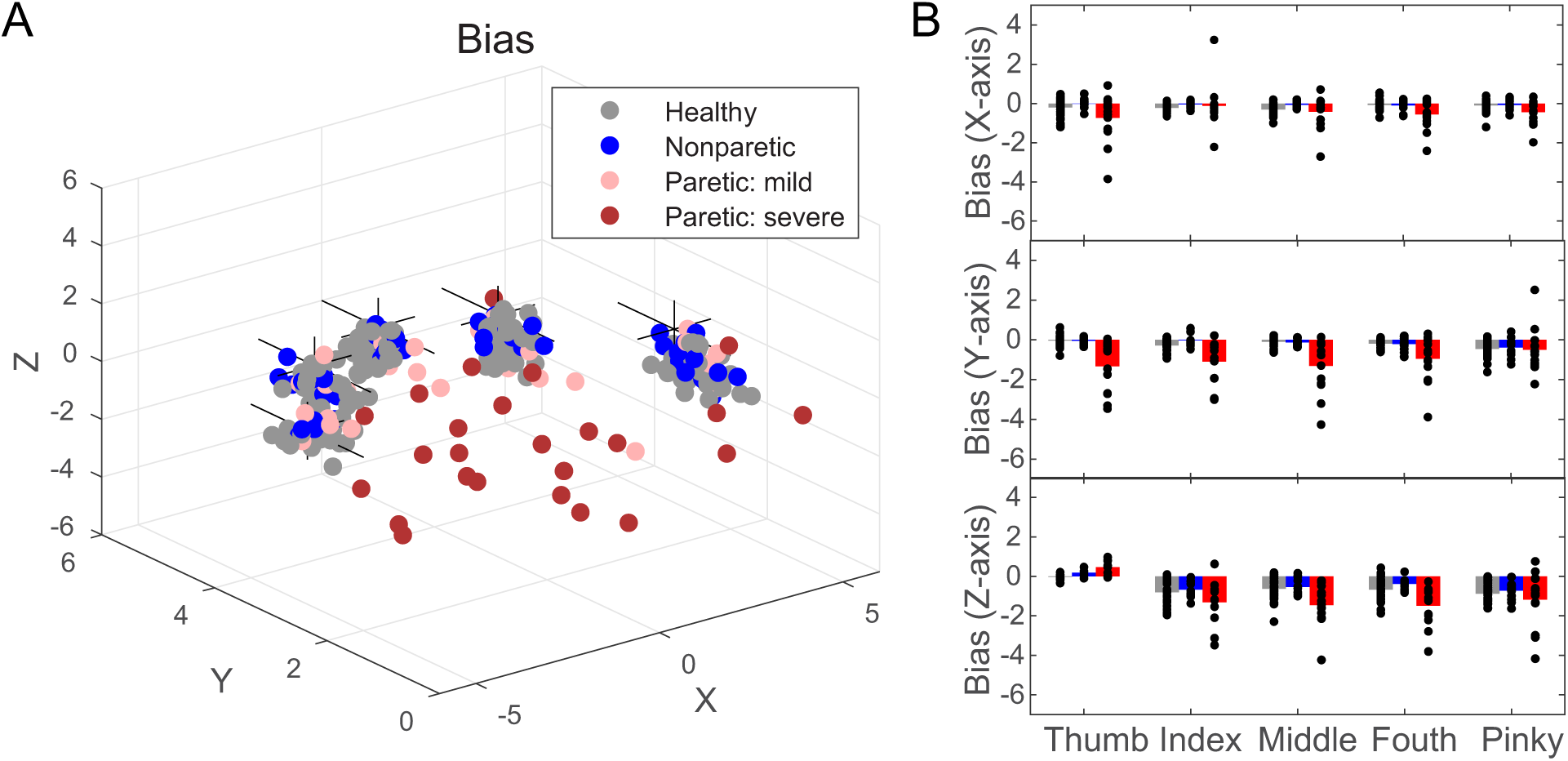
Bias measures. **A.** Mean biases for each subject for all fingers plotted in the same 3D space. Paretic data are separated by severity: (FMA>=40 vs. (FMA<40)). **B.** Bias summary in the flexion (-Y and -Z), extension (+Y and +Z), and abduction/adduction (+X/-X) directions for the three groups.

We first tried to tease apart the contributions of subcortical versus biomechanical sources to enslaving patterns. To examine if the two measures for lower-level constraints share the same sources, we first tested the predictability of resting-hand postures (Mount Distance and Mount Angle) to Bias in both the healthy and paretic hands. Intriguingly, for all groups, these resting-posture measures were not significant factors for biases (in non-pathological hands: Mount Distance *p* = 0.50, Mount Angle *p* = 0.096; in paretic hands: Mount Distance *p* = 0.32, Mount Angle *p* = 0.92). This suggests that biases shown in voluntary finger control may not share the same skeletal and tendentious structural constraints as resting hand postures. We then tested the effect of postures on enslaving patterns. Similarly, resting-hand posture variables were not significant predictors for enslavement/coactivation patterns in all groups (non-pathological: Mount Angle *p* = 0.31, Mount Distance *p* = 0.33; paretic: Mount Angle *p* = 0.79, Mount Distance *p* = 0.89). We further explored this question by looking at biases in the non-paretic hand (in stroke participants only). A similar LME model was used with the Bias values in the paretic hand replaced by those from the non-paretic hand. Interestingly, the Non-paretic Bias was not a significant predictor of either Bias (*p* = 0.63) or Enslavement in the paretic hand (*p* = 0.29). This result thus ruled out the contribution of biomechanical factors entirely.

We then conducted LME analyses to test our main hypothesis that low-level biases may be able to account for finger enslavement patterns in the paretic hand, but not the coactivations in the non-pathological hands. Variables submitted to the LME model were Instructed Finger, Target Direction, Group (healthy, non-paretic, paretic), Enslaved Finger and Direction (direction of uninstructed finger activity), Bias, Mount Angle, and Mount Distance. The modeling results showed that Bias was a strong predictor of enslavement patterns in the paretic hands (*p* < 2e-16), but not in the non-pathological hands (*p* = 0.17). Additionally, Enslaved Direction (*p* = 0.02) and Enslaved Finger (*p* = 2.94e-07) were both significant predictors for paretic enslaving patterns, indicating these patterns are finger and direction specific.

The LME analysis further confirmed that coactivation patterns in the healthy hand were mainly driven by top-down task goals: Instructed Finger (*p* = 5.57e-05), Target Direction (*p* < 2e-16), and Enslaved Direction (*p* = 8.21e-14) were all significant predictors.

An increase of flexor bias has been viewed as an intrusion due to neural upregulation of the subcortical pathways (Zaaimi et al. 2012; Choudhury et al. 2019). As all groups showed a flexor bias, with the paretic hand presenting a stronger version, we thus further teased apart the normal vs. intruding subcortical contributions to enslavement patterns in the paretic hand. To isolate flexor biases, we used the mean of Biases in the Y and Z directions for each finger, and then used this Flexor Bias as a predictor for the Individuation Index in LME analysis. Individuation Index is an overall net sum of individuation impairment after stroke, so if the flexor bias dominates the entire hand, regardless of which finger the participant intended to move, we should expect that flexor bias would be a strong predictor for the Individuation Index in non-flexion directions, ab/adduction (X axes), but not the instructed finger. Indeed, in the non-pathological hand, while Instructed Finger was a strong predictor (*p* = 1.83-08), Flexor Bias was not (*p* = 0.06); in contrast, Flexor Bias was the single significant predictor for the Individuation Index in ab/adduction directions in the paretic hand (*p* = 7.48e-05), but Instructed Finger was not significant (*p* = 0.39).

### Loss of complexity and intrusion of flexor bias are dissociable in the paretic hand

The above analyses ruled out the contribution of biomechanical constraints and suggested that the intrusion of flexor bias might be the key player in finger individuation impairment. We then moved on to examine if the two other factors, loss of complexity and the intrusion of flexor bias: do both contribute to the observed enslaving patterns in the paretic hand, or one of them dominates the impairment? Further, if both are contributing factors, to what extent are they dissociable? To answer these questions, we tested the predictability of flexor intrusion on Angular and Euclidean Distances of the paretic hand enslavement. Given the above converging evidence that Angular Distance is a measure of complexity of geometric shape of finger coactivation, whereas Euclidean Distance is more sensitive to the magnitude change of these patterns across task goals, if the two factors are dissociable, the intrusion of flexor bias would predict the *magnitude* (Euclidean Distances), but not the *shape* (Angular Distances) of the enslavement patterns.

LME models with fixed factors of Finger and Bias and a random factor of Subject showed that while Bias was not a significant predictor for Angular Distances (distances by Instructed Finger: *p* = 0.08; by Target Direction: *p* = 0.30), it was a significant predictor for the Euclidean Distances (distances by Instructed Finger: *p* = 9.24e-05; by Target Direction: *p* = 1.70e-11) in non-pathological hands. Similar results also hold with the paretic hand: Bias was not a significant predictor of Cosine Distance by Instructed Finger (*p* = 0.47) and a weak predictor by Target Direction (*p* = 0.03), whereas it was a strong predictor for Euclidean Distance (by Finger: *p* = 7.98e-06; by Target Direction: *p* = 0.009). It’s intriguing that Bias could weakly predict Cosine Distance by Target Direction, i.e., pattern distances across conditions where different fingers moving towards the same target direction could be partially explained by flexor biases. As shown in Fig. 5 C-D, a large reduction of Angular Distances in severe patients was shown in -X, -Y, and -Z directions, which coincides with the directions that may easily induce flexor bias.

While the fact that bias could predict Euclidean Distances in both non-pathological and paretic hands suggests that sub-cortical factors may be the major contributor to the magnitude of enslaving patterns, the above results did not directly speak to the cause of the *change* of the magnitude in enslaving patterns due to stroke. To closely examine the contribution of the *intrusion* of flexor bias to Angular vs. Euclidean Distances in the paretic hand, we first computed the difference between paretic - non-paretic Bias as an estimate of intrusion of flexor bias (Bias Difference), and then used it as a predictor in the LME models for the two types of distances in the paretic and non-paretic hands. Indeed, Bias Difference was not a significant predictor of Angular Distance in both paretic and non-paretic hands (non-paretic by Finger: *p* = 0.88, by Direction: *p* = 0.66; paretic by Finger: *p* = 0.20, by Direction: *p* = 0.07), indicating that intrusion of flexor bias could not fully account for the loss of complexity.

In contrast, Bias Difference was a highly significant predictor of Euclidean Distances in the paretic hand (by Finger: *p* = 1.08e-07; by Direction: *p* = 0.008) (Fig. 9). Intriguingly, while Bias Difference was not a significant predictor for the Euclidean Distances in the non-paretic hand by Finger (*p* = 0.29), it was significant by Direction (*p* = 0.0009). Post-hoc correlation analysis indicated that Bias Difference and Euclidean Distances were negatively correlated in the non-paretic hand (Fig. 9D), suggesting that the magnitude of coactivation patterns in the non-paretic hand was following the opposite trend of intrusion of flexor biases. This further corroborates the above results indicating that Euclidean Distance between different enslaving patterns across task goals is mainly driven by lower-level biases. Together, the above results confirmed our hypothesis that loss of finger control complexity and intrusion of flexor bias are distinct processes contributing to the impairment in finger control after stroke.

**Figure 9.**
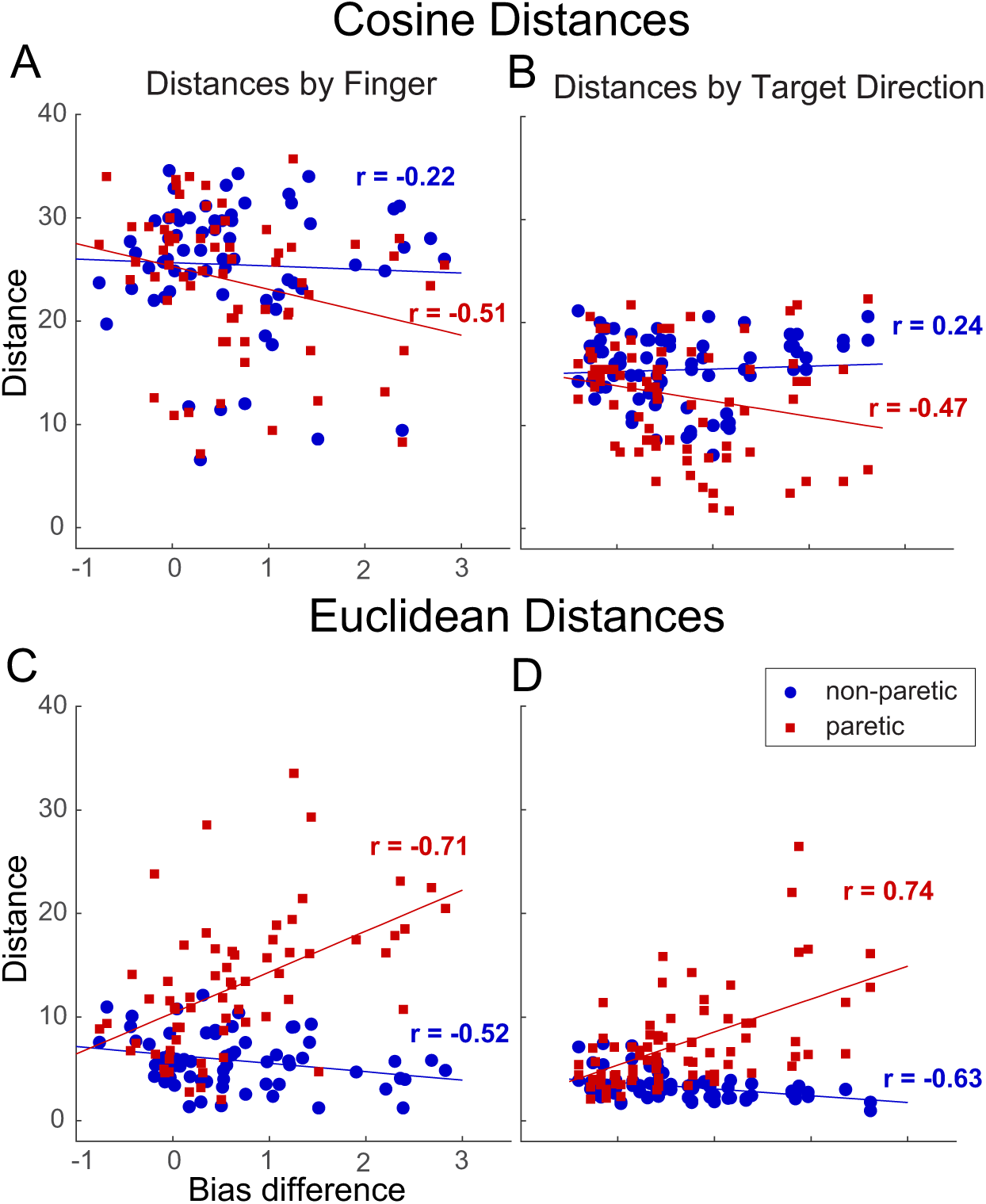
Scatter plots of Bias Difference between paretic and non-paretic (intrusion of flexor bias) and distance measures. **A-B.** Cosine Distances by Finger and Target Direction, respectively, and **C-D.** Euclidean Distances by Finger and Target Direction, respectively. r-values are Pearson correlation coefficients.

### Finger coactivation patterns in the healthy hand require a higher number of PCs to account for than the enslavement patterns in the paretic hand

Lastly, to test the idea that the human nervous system uses a small number of synergies in complex manipulations (Overduin et al. 2012; Santello and Soechting 1998; Latash 2009), we also used a data-driven approach to analyze the finger coactivation/enslavement patterns by PCA. Specifically, we tested the hypothesis that a small set of PCs can account for a large amount of variance in the finger coactivation/enslavement patterns observed in the 3D Individuation task. If the patterns in the paretic hand present a similar but exaggerated pattern as those in the healthy hand, we would expect a similar number of PCs to account for finger coactivation/enslavement patterns in healthy and paretic hands. Instead, our results showed that to achieve the same level of variance explained, a larger number of PCs will be needed for the healthy and non-paretic hands compared to the paretic hands (Fig. 10). This finding is also consistent with the idea that the paretic hand loses the complexity of the hand usage that could not be explained by a handful of PCs present in all hand usage (Yan et al. 2020).

**Figure 10.**
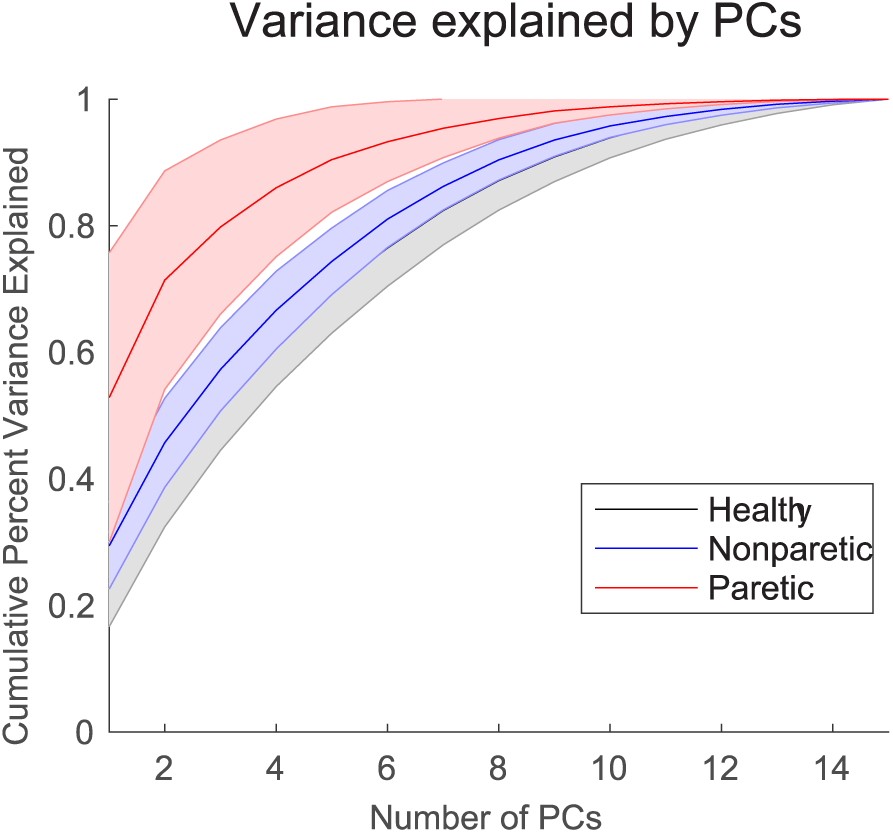
PCA analysis results. Variance explained by the number of PCs in finger coactivation/enslavement patterns for the healthy, non-paretic, and paretic hands. Note that there are a maximum of 15 PCs due to the nature of the individuation task.

## Discussion

In this study, we investigated finger individuation of the whole hand in all three dimensions to examine three potential factors that may contribute to the loss of individuated finger control after stroke: loss of complexity, intrusion of flexor bias, and biomechanical constraints. Our results demonstrated that finger enslavement patterns recorded from 3D isometric finger forces during a task that assessed the full capacity of finger individuation after stroke were qualitatively different from the coactivation patterns observed from a healthy hand. The patterns in a healthy hand were mainly driven by top-down control (e.g., task goals such as instructed finger and target direction), with the patterns differing by shape when task goals change. In contrast, enslavement patterns observed in the paretic hand present reduced variances in shape across task goals and an increased magnitude, which could largely be accounted for by lower-level biases. In addition, biomechanical factors assessed by resting hand posture could not account for both the biases and enslavement in the paretic hand. Importantly, the intrusion of flexor biases, which could not account for the reduced complexity in enslavement patterns in the stroke hand, was the single factor that predicted the magnitude of the pattern changes across task goals. These findings depart fundamentally from the previously established view that enslavement after stroke is an exaggerated version of those found in the healthy hand (Z. M. Li, Latash, and Zatsiorsky 1998; S. Li et al. 2003). Instead, our results suggest loss of finger individuation after stroke may be due to the interplay of two distinct sources: a loss of higher-level cortical controls that produce complex finger- and direction-specific patterns in the healthy hand, and an intrusion of flexor bias, possibly due to an up-regulation of subcortical control.

### Finger coactivation patterns as a window to the complexity in hand control

Complexity in healthy hand control has been examined in the past (e.g., (Yan et al. 2020; Todorov and Ghahramani 2004)). However, the assessment tools used in those studies, e.g., the CyberGlove, had an unsatisfying resolution that cannot meet the level of dexterity humans use their fingers with; participants were often engaged in unconstrained object manipulation tasks, which, despite its own advantages, lacks the careful experimental control to tease apart underlying neural control processes. To our knowledge, our study is the first to carefully quantify finger coactivation patterns in the healthy and stroke hands in 3D and clearly tease apart factors contributing to the enslaving patterns observed in stroke: loss of complexity vs. lower constraints set by biomechanical and subcortical factors, considering both healthy baseline and pathological upregulated forms. While here we used Instructed Finger and Task Direction to proxy top-down control factors, the same phenomenon may be generalizable to more complex manual tasks, such as object manipulation.

Most previous complex finger control studies were examining finger kinematics. Here we used a sensitive device to record isometric fingertip forces. We render forces detected from all five fingertips and project them in a 3D virtual task space. It has been shown that motor control principles in isometric force production may be generalizable to limb position control (Rotella et al. 2014). We are thus confident that findings using the current 3D isometric force recording will hold in kinematic spaces. Moreover, the high sensitivity of our device and the task environment allows us to capture the complexity of finger control more faithfully.

### Neural control mechanisms for the complexity of finger control

Our results, showing the task-dependent nature of healthy hand finger coactivation patterns, are in line with recent findings that hand posture in higher dimensions (low-variance PCs) is dependent on both the hand structure and behavior tasks (Yan et al. 2020). The authors of that study suggested that those low-variance PCs in hand postures during complex tasks were not simply motor noise, but may participate in hand movement repertoire in meaningful ways. Other researchers have also suggested that fingertip force fluctuations reveal flexible rather than synergistic controls (Kutch et al. 2008). These coactivation patterns in the task-irrelevant fingers in a healthy hand may play an important functional role in complex hand movement. One possibility is that they help to optimally balance the efficiency and precision of the fingers directly engaged in the task. Indeed, when the hand is under the pressure of improvising different finger positions and trajectories de novo while maintaining steady and accommodating hand postures during a complex manual task, a fixed pattern of finger coactivation would be conceivably suboptimal.

Our results show that paretic hands, especially those with moderate-severe impairment, lose complexity in finger control, i.e., reduced directional differences across different task goals. Our previous study has demonstrated that CST damage is strongly correlated with finger individuation impairment and recovery (Xu et al. 2017). The loss of complexity observed in the current study may share the same neural mechanisms. Fine finger control has long been suggested to stipulate a unique evolutionarily new descending pathway, namely the monosynaptic corticomotorneuronal (CM) projections within the CST, projecting from the anterior bank of the central sulcus to spinal motor neurons (Lemon 1993; Rathelot and Strick 2009). Loss of fractionated finger control has been especially associated with the disruption of this pathway (Lawrence and Kuypers 1968b; 1968a). Besides the CM cells directly influence spinal motoneurons via monosynaptic projections (Fetz, Cheney, and German 1976; Lemon, Mantel, and Muir 1986), other indirect descending projections include spinal segmental interneurons, C3-C4 propriospinal neurons, pontomedullary reticular formation, and red nucleus (Kuypers 1987). It has been suggested in the literature that the divergent CST inputs may play an essential role in sculpting the neural output driven by subcortical pathways, e.g., the reticulospinal tract (RST) to increase the flexibility of motor repertoire and achieve more precise finger control (Schieber 1990; Schepens and Drew 2006; Riddle and Baker 2010). In addition, intracortical activities may also play a role in shaping these coactivation patterns (Sohn and Hallett 2004; Beck and Hallett 2011). We thus speculate that the healthy task-driven complex patterns in finger coactivation may be a result of an intricate balance of the cortical and subcortical interplay.

### Biomechanical constraints do not contribute to enslavement in stroke

Human hands are evolved from gross prehension, where all five fingers open or close together to grasp an object. It is thus not surprising that we observe a strong flexor bias in all participants, from healthy to non-paretic to paretic hands. Multiple sources may contribute to this bias, from biomechanical, such as skeletal and muscular structures and connecting tissues, to last-order neuronal input, to various subcortical systems, such as the reticulospinal and rubrospinal pathways. Intriguingly, in the current study, we found that resting hand posture could explain neither the biases in the instructed fingers, nor the non-instructed finger enslaving patterns in any group. This result ruled out the possibility of skeletal and tendentious structural contributions to the impairment of finger individuation. Further, within-subject comparisons showed that biases in the non-paretic hand could not predict enslaving patterns in the paretic hand either. Although both biomechanical and subcortical factors may contribute to the non-pathological hand biases, this finding, together with the resting-hand-posture finding, completely ruled out contributions from the biomechanical factors. Similar ideas have been suggested in the previous literature: while finger coactivation during passive movement can be largely attributed to biomechanical constraints, changes in those patterns in voluntary finger control after stroke, especially during isometric force tasks, have been found to be largely due to neural sources (Lang & Schieber, 2004, Li et al. 2003, Kamper et al. (2003)). Other researchers also pointed out that synergies may not necessarily arise from biomechanical constraints, but as a result of optimal control solutions (Diedrichsen, Shadmehr, and Ivry 2010).

### Loss of complexity and intrusion of flexor biases after stroke may arise from distinct biological processes

While the flexor bias in the non-paretic hand could not predict the enslaving patterns in the paretic hand, we found that those in the paretic hand were a strong predictor for its own enslaving patterns. Strikingly, biases along the flexion/extension directions (Y and Z directions in the Cartesian space) in the instructed fingers strongly predicted individuation impairment in ab/adduction. This finding already suggests that it is the extra amount of flexor bias (or the intrusion of flexor bias), instead of a normal form of flexor bias, that may account for the enslaving patterns observed in the paretic hand.

Consistent with the literature (Kline, Schmit, and Kamper 2007), we found that paretic hands present a stronger version of flexor bias compared to the non-paretic hand. Here we used the difference of biases across the paretic and non-paretic hands as an estimate of the intrusion of flexor bias. While one may think that since biases seem to dominate the enslaving patterns and individuation impairment of the entire hand, it may be the single factor that accounts for everything observed in a paretic hand, this does not seem to be the case. Our measure for the intrusion of biases could not predict the loss of complexity in enslaving patterns, i.e., angular distances in the paretic hand.

The paretic hand enslaving patterns exhibited not only a reduction of geometric shape differences (angular distances), but also an increased magnitude of distances (Euclidean distances) across task goals. The more severe the impairment, the more reduction of angular distances and more increase of Euclidean distances, suggesting that these two distance measures assess distinct aspects of individuation impairment. The key question is if these changes in enslaving patterns assessed by these two measures share the same underlying mechanism. Our analyses using the intrusion of biases appear to support the dissociation of the two: while the intrusion of flexor biases was not a significant predictor for angular distance in both paretic and non-paretic hands, it was a strong predictor for Euclidean distance in the paretic hand. These findings suggest that the two manifestations of finger control impairment appear to share different sources, with the former a peeling of finger control complexity while the latter overwhelmed by an exaggerated flexor bias. This finding is intriguing but also consistent with biological findings of distinct descending pathways and biological recovery processes contributing to motor impairment after stroke.

A parallel question is thus what the neural origins for the three types of behaviors are: complexity, biases in the healthy hand, and those in the paretic hand. As mentioned above, complexity is mainly driven by the top-down CST input. The finding that enslavement and biases in the paretic hand cannot be simply accounted for by the biases in the non-paretic hand suggests that these two kinds of flexor biases may not be due to the same biological origin. It has been suggested in human studies that the intrusion of flexor biases may be due to upregulations of subcortical pathways, such as reticulospinal tract (Kamper et al. 2003; Ellis et al. 2012; Choudhury et al. 2019). Additionally, evidence of regeneration and strengthening of the reticulospinal tract participation in hand function recovery after lesions of CST has been found in non-human primate studies (Zaaimi et al. 2012). Particularly, the strengthened connections were found to be projecting to flexors, not extensors, which may explain the intrusion of flexor biases. The role of RST in hand function, however, is limited due to its non-specific innervation to muscles. Thus, the newly strengthened pathways may not be able to compensate for the loss of complexity due to lesions in CST. They may set a constraint to finger individuation recovery and could even interfere with it by adding the intrusion of a strong flexor bias due to its imbalanced projections. Together, the intricate interplay between the reduction of cortical input and subcortical upregulation may explain the behavioral dissociation between the loss of complexity and intrusion of flexor bias we observed in the enslaving patterns after stroke.

## Conclusions

In a cohort of chronic stroke patients, we tested the contributions of biomechanical, subcortical, and cortical input to the observed impairment of finger individuation in the paretic hand, using a sensitive 3D isometric force device with an individuation task, and a set of comprehensive pattern analyses. Our results not only ruled out the contributions of biomechanical factors, but also the baseline level of flexor biases in a normal hand, to the loss of individuated finger control. We further demonstrated that the loss of complexity and intrusion of flexor biases have dissociable factors combined to produce the enslaving patterns observed in our 3D finger individuation task. These findings show that loss of individuated finger control after stroke may be a result of two distinct sources: a loss of higher-level cortical controls that produce complex finger- and direction-specific patterns in the healthy hand, and an intrusion of flexor biases possibly due to an upregulation of subcortical control. Together, our findings challenge a fundamental assumption prevailing in the field that enslavement patterns after stroke are an exaggerated version of healthy hand coactivation patterns. Together, our sensitive device, assessment paradigm, and analysis methodology, as a package, also demonstrate an effective approach to assessing hand control behavior using finger-coactivation patterns to infer distinct underlying biological sources.

## Materials and Methods

### Participants

Twenty-nine healthy participants (age range 19-43, 16 women, all right-handed (Oldfield 1971)) and thirteen stroke patients (age range 30-80, 7 women, 12 right-handed) participated in the study. Nine patients were affected on the left side. All patients met the following inclusion criteria: (1) age 21 years and older; (2) ischemic stroke at least 6 months prior; (SEM), confirmed by computed tomography, magnetic resonance imaging, or neurological report; (3) residual unilateral upper extremity weakness; (4) ability to give informed consent and understand the tasks involved. We excluded participants with one or more of the following criteria: (1) upper extremity Fugl-Meyer assessment (FMA) >63/66 (Fugl-Meyer et al. 1975), (2) cognitive impairment, as seen by a score of <20/30 on the Montreal Cognitive Assessment (MoCA); (2) history of a physical or neurological condition that interferes with study procedures or assessment of motor function (e.g., severe arthritis, severe neuropathy, Parkinson’s disease, space-occupying hemorrhagic transformation, bihemispheric stroke, traumatic brain injury, encephalopathy due to major nonstroke medical illness, global inattention, large visual field cut (greater than quadrantanopia), receptive aphasia (inability to follow 3-step commands), inability to give informed consent, major neurological or psychiatric illness); (3) inability to sit in a chair and perform upper limb exercises for one hour at a time; (4) terminal illness. For detailed patient characteristics, see Table 1. All participants signed a written consent, and all procedures were approved by Institutional Research Board at Johns Hopkins University.

### The Apparatus and 3D Force Recording

#### The HAND

To fully test people’s finger control abilities in all directions, we have designed a new device, the Hand Articulation Neuro-training Device (HAND, JHU reference #C14603) that can detect micro-isometric forces at the fingertips in 3D (Fig. 1A). A custom-developed highly sensitive finger-force sensor using strain gages (at the milli-Newton level), is built underneath each fingertip. All data were digitized by a custom-built data acquisition (DAQ) unit, and sampled at 1KHz per strain-gauge-pair comparison. The streamed and digitized force data were projected on the computer screen in real-time during the task, with a scaling of 0.6 cm/N, 0.4 cm/N, and 0.8 cm/N in the X, Y, and Z directions in a virtual 3D space, respectively. All behavioral tasks in the virtual space were programmed using custom Python scripts. (van Rossum and Drake 2011).

#### Setting up the hand

Participants were seated comfortably at a desk with their tested forearms resting on the table, their hands resting in a neutral posture with wrist pronated, and all five fingers fitted in the soft custom-fit silicon cups (Fig. 1B). The position of each finger could be adjusted to achieve the most relaxed hand posture. During setup, forces produced at each fingertip were constantly monitored on the computer screen as vertical bars to find the posture that minimized forces (i.e., <1 N) produced by all fingers (Fig. 1C). Once the minimum forces were achieved, the hand posture was fixed throughout the entire experiment. The mounting angle and distance for each finger (Fig. 1B) were recorded as the resting hand posture.

### Experimental Design and Procedures

#### Natural finger trajectories

After fitting the hand in the device, participants were instructed to move a dot in the virtual 3D space on the computer screen (Fig. 1E) by exerting an isolated isometric force (max = 10 N) with one finger-joint in the instructed direction repeatedly for 10 seconds (30 seconds for stroke patients). The goal of this step was to map out the natural force trajectories in 3D space for each finger joint in isolation. Movement of the dot along the virtual xyz axes reflected forces exerted by MCP ab/adduction, PIP flexion/extension, and MCP flexion/extension, respectively (Fig. 1D).

#### Finger-individuation task

After natural trajectories for each finger were recorded, the mean trajectories in the six directions were computed by taking the first eigenvector of the overall trajectories in each direction. Four force targets were defined along this mean vector for each of the six directions at 20, 40, 60, and 80% of the maximum extent (Fig. 1E). After computing the natural trajectories, participants completed the individuation task by producing an isolated isometric force with a single finger joint to hit one of the four targets defined along the finger’s natural path. Critically, participants were instructed to keep the other fingers immobile while exerting the isolated force with the specified finger. The overall force levels in the uninstructed fingers were monitored by a vertical bar on the computer screen (Fig. 2A&C). The goal is to keep the bar as low as possible while hitting the target. Once reached, the target would disappear, and the participant was instructed to relax the active finger back to the home position. An important feature, and advantage, of our HAND device is that the virtual space can be calibrated based on the patient’s maximum force level, which allows very weak patients with near-plegic hands to see clear feedback on their finger activities. Four trials per finger per target were tested, amounting to a total of 480 trials per hand. All patients were tested with both the affected and unaffected hands, and all healthy participants were tested with their dominant hands.

### Clinical Assessments

To compare with other clinical studies, all patients were also assessed with clinical impairment and functional measures during their visit. For clinical impairment assessment, we tested upper-extremity motor impairment using the Fugl-Meyer Upper-extremity Assessment (FMA) (Fugl-Meyer et al. 1975), grip strength using dynamometry, and sensory impairment using the Semmes-Weinstein Monofilament (Bell-Krotoski and Tomancik 1987) assessment. For functional assessment, we tested patients’ hand function using the Action Research Arm Test (ARAT) (Lyle 1981).

### Data Analyses

#### Force data preprocessing

All sampled force data from each trial was smoothed using a second-order low-pass filter with a cutoff frequency of 5Hz, converted to Newtons, and normalized by the baseline (mean of the first 100 data samples). To make data consistent and easy to interpret across fingers, the thumb data was rotated 90 degrees in the Y and then Z planes in data preprocessing, such that the thumb orientation would be the same as the four fingers in a pronated posture.

#### Individuation Index

We expand upon our previously established method to calculate the Individuation Index (Xu et al. 2017) to assess participants’ overall ability to individuate each finger in 3D. Uninstructed-finger activations were computed as the mean deviation (*meanDevP*) from baseline forces,

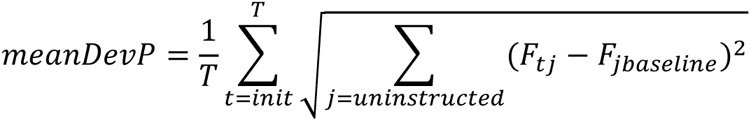

where the root mean square of all uninstructed finger forces from the baseline were summarized over the instructed finger activation period, which is defined from the initiation of active force (> 0.02 N) to the minimum force velocity after the peak (Fig. 3A). The Individuation Index is then calculated as the regression slope of the overall meanDevP from all uninstructed fingers against active finger force levels. One Individuation Index is computed for each instructed finger at each target direction (X, Y, Z) that the finger attempts to move to.

#### Enslavement/coactivation

Similar to biases measures, enslavement/coactivation is calculated as log ratios of forces in an uninstructed finger/direction with respect to an instructed finger or target direction. For example, the enslavement of finger *f_i_* along the x-axis with respect to finger *f_j_* instructed to move towards positive x would be:

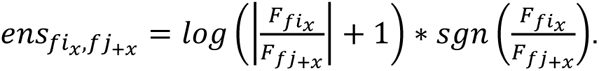

To account for the fact that both instructed and uninstructed fingers can move toward positive or negative directions, the log transformation was applied to the absolute values of the ratio, a scalar 1 was added to the ratio, and the original sign was then put back in to preserve the direction of coactivation patterns.

#### Biases measure

To capture the lower-level factors that may contribute to finger coactivation patterns, we derived two bias measures: 1) Trajectory biases were derived from force recordings in the instructed fingers: Biases *along* each axis are computed as the log ratio of forces produced in the positive over the negative direction for that axis,

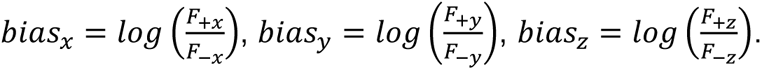

This measure captures a finger’s “preferred” direction in XYZ axes in voluntary action (Fig. 8A). 2) We also record the resting hand posture after each hand was fit in the HAND device (see The Apparatus and 3D Force Recording) as the hand’s physical biases.

#### Reliability analysis

To determine the reliability of our major measures, Individuation Index, Enslavement, and Bias, we calculated the split-half reliability. To account for the increased variability when splitting data in halves, the formula 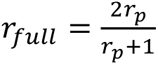 was used.

#### Pattern similarity analyses of finger coactivation/enslavement

To directly test the hypothesis that finger coactivation patterns in the healthy hand are mainly driven by top-down influences (i.e., task demand), we directly compared the patterns across different instructed fingers and target directions using Representation Similarity Analysis (RSA). Representation distance matrices (RDM) (Fig. 4 A-B & D-E) for each instructed finger were constructed by computing the angular (Cosine) distance between pairs of uninstructed-finger coactivation patterns (log ratios, see above) when the same instructed finger attempted to move to two different target directions. To directly compare coactivation/enslavement patterns for each instructed finger, we first arranged the finger coactivation matrices (4 uninstructed fingers X 3 directions) per-finger per-target direction. An angular (Cosine) distance is computed between the same uninstructed finger across two different target directions. This step will result in a 4×1 array. The average across all four uninstructed fingers was then placed in the instructed finger representation distance matrix (RDM). For example, when the index finger moves, the uninstructed fingers would be the other four fingers: thumb, middle, fourth, and little fingers. Distances of the forces in these four fingers between the index finger moving toward two different directions (e.g., +X vs. +Y) were computed in a pairwise manner, and summed to one distance value in the RDM for the index finger. (Fig. 4A). This step will result in five RDMs for each instructed finger. The same procedure was taken to compute the RDMs for six target directions to directly compare the pattern similarities across instructed fingers. To compare the shape vs. magnitude of coactivation/enslavement patterns, we also used Euclidean distances to assess the magnitude of pattern differences.

#### Linear mixed-effect model analyses

All statistical analyses were performed using custom-written MATLAB and R (Team, n.d.) routines. To test the group differences in Individuation Indices, as well as the role of top-down (Instructed Finger and Target Direction) vs. low-level factors (Bias, Drift, and Hand Posture (Mounting Angle and Distance)) in enslavement/coactivation patterns, we used linear mixed-effect (LME) models implemented in the *lme4* package (Bates et al. 2015) in R. In these models, Participant was treated as a random effect, and other aforementioned factors were treated as fixed effects. Statistical significance of fixed factors and their interactions were tested using the likelihood ratio test using *X*^2^ distribution.

#### Synergy analyses

We performed a Principal Component Analysis (PCA) on the finger forces from all fingers, across all conditions: instructed fingers and target directions for each group of subjects. Finger forces used in this analysis were those recorded from all fingers at the time when the instructed finger reached the target. To examine variance explained by the number of PCs, we normalized the eigenvalues by their sum to obtain the fraction of variance explained by each eigenvector. A cumulative sum was taken of these values to get the fraction of variance explained by the first *n* components. The progression of the number of components and variance explained were compared across three groups: paretic, non-paretic, and healthy hands.

## Acknowledgements

We thank Dr. Alkis Hadjiosoif for assistance in stroke patient recruitment, and Aiden Devaney and Teresa George for assistance in sensor calibration and validation. This work was supported by Johns Hopkins Malone Center Seed Grant 80046364 (JX), NIH CTSA UL1 award 2UL1TR002378-06 (JX). The authors declare no conflict of interest.

